# Live Imaging of Epithelial Phagocyte Differentiation in the *Drosophila* Ovary Reveals Transient and Novel Behaviors

**DOI:** 10.64898/2026.05.21.726826

**Authors:** Caique Almeida Machado Costa, Cristian Santiago Santiago, Jie Sun, Yi-Chun Huang, Wu-Min Deng

## Abstract

The transformation of epithelial cells into Non-Professional Phagocytes (NPPs) is a conserved and versatile adaptation that occurs in response to immune challenges, tissue remodeling, and apoptotic debris clearance. In the *Drosophila melanogaster* ovary, follicle cells (FCs) acquire phagocytic capabilities under stress, providing a powerful model to study this process. Using time-lapse live imaging, we captured dynamic behaviors associated with FC-to-NPP differentiation that are too transient to detect by static imaging. Our approach confirmed established features, including germline cell death, cytoplasmic expansion, and debris engulfment, and revealed previously unrecognized capabilities. These include a gradual increase in JNK pathway activation, after which NPPs exhibit collective migration toward dying germline cells, epithelial delamination, long-range target capture through pseudopodial extensions, and the engulfment of neighboring FCs. These findings demonstrate that epithelial-derived NPPs can perform complex phagocytic tasks typically attributed to professional phagocytes such as macrophages. Our work establishes the *Drosophila* ovary as a robust *in vivo* system to uncover conserved and novel aspects of epithelial plasticity and phagocytic function, particularly those involving transient behaviors missed by fixed-sample analyses.

## INTRODUCTION

Phagocytosis is a dynamic cellular process essential for the elimination of pathogens and the clearance of apoptotic debris (Uribe-Querol and Rosales, 2020). This activity typically requires cells to undergo distinct morphological transformations that facilitate tasks such as migration, cytoplasmic expansion, and ultimately the engulfment of dying cells (Mylvaganam et al., 2021). While professional phagocytes such as macrophages are specialized for this function, epithelial cells can, under specific circumstances, particularly in the presence of nearby dying cells, acquire phagocytic capabilities and function as NPPs (Arandjelovic and Ravichandran, 2015; Monks et al., 2008). This phenomenon has been documented across the animal kingdom, including in mammals, where NPPs have been observed in diverse tissues such as the lungs, vasculature, skin, myocardial tissue, and brain (Seeberg et al., 2019; Sihombing et al., 2021).

*Drosophila melanogaster* serves as a powerful model system to study this transition. In the fly ovary, developing egg chambers undergo successive developmental stages to fully differentiate into an egg ready to be laid. These egg chambers consist of germline cells surrounded by a monolayer of epithelial-FCs (Jia et al., 2016). Under stress conditions such as radiation (Panagopoulos et al., 2007), nutrient deprivation (Serizier and McCall, 2017), warm acclimation (Gandara and Drummond-Barbosa, 2022), cold acclimation (Lirakis et al., 2018), or persistent activation of Notch signaling in follicle cells (Costa et al., 2025; Huang et al., 2025) germline cells within mid-stage egg chambers (Stages 7-9) can undergo regulated cell death to halt the energetically costly process of oogenesis. In response, adjacent FCs adopt NPP behavior and phagocytose germ cell corpses.

This process has been characterized as a coordinated program of cellular and metabolic remodeling that involves activation of signaling pathways, phagocytic receptors, autophagy, cytoskeletal reorganization, and endolysosomal trafficking (Barth et al., 2011; Costa et al., 2025; Huang et al., 2025; Lebo et al., 2021; Serizier et al., 2022; Serizier and McCall, 2017). While live imaging studies have demonstrated recruitment of extracellular acidification machinery by NPPs during germline clearance in late oogenesis (Mondragon et al., 2019), the mechanisms by which phagocytic capabilities are initiated and coordinated during stress-induced <u>g</u>ermline <u>d</u>eath <u>a</u>nd <u>c</u>learance (GDAC) at early stages of oogenesis remain poorly understood. To address this gap, we focused on mid-stage egg chambers and employed time-lapse live imaging to capture transient cellular behaviors that cannot be resolved using static imaging approaches.

Here, we show that mid-stage ovarian FCs progressively acquire coordinated migratory and phagocytic behaviors during stress-induced differentiation into NPPs. Live imaging revealed gradual emergence of epithelial delamination, long-range debris capture via pseudopodial extensions, and the ability to engulf not only dying germline cells but also neighboring FCs. Overall, these findings define NPP differentiation as a dynamic cellular state transition that integrates epithelial plasticity with immune-like effector functions.

## RESULTS

### *Ex vivo* live culture and imaging reveal germline cell death and FC differentiation into NPPs

We began our analysis by asking whether live culture conditions are sufficient to induce germline cell death and drive FC differentiation into NPPs, similar to the well-established protocol in which flies are exposed to 16–20 hours of nutrient deprivation (Lebo et al., 2021). To test this, we cultured *wild-type* (*w¹¹¹⁸*) *Drosophila* ovaries in *ex vivo* culture medium for 18 hours (Fig. 1A1). As controls, we used ovaries dissected from well-fed flies and from flies subjected to 18 hours of starvation (Fig. 1A2). Comparable to the starvation condition, live culture increased the frequency of germline cell death in mid-stage egg chambers (Fig. 1B), as evidenced by staining for cleaved (active) Dcp-1 (Song et al., 1997) in germline cells of egg chambers (Fig. 1C2, C4).

**Figure 1.**
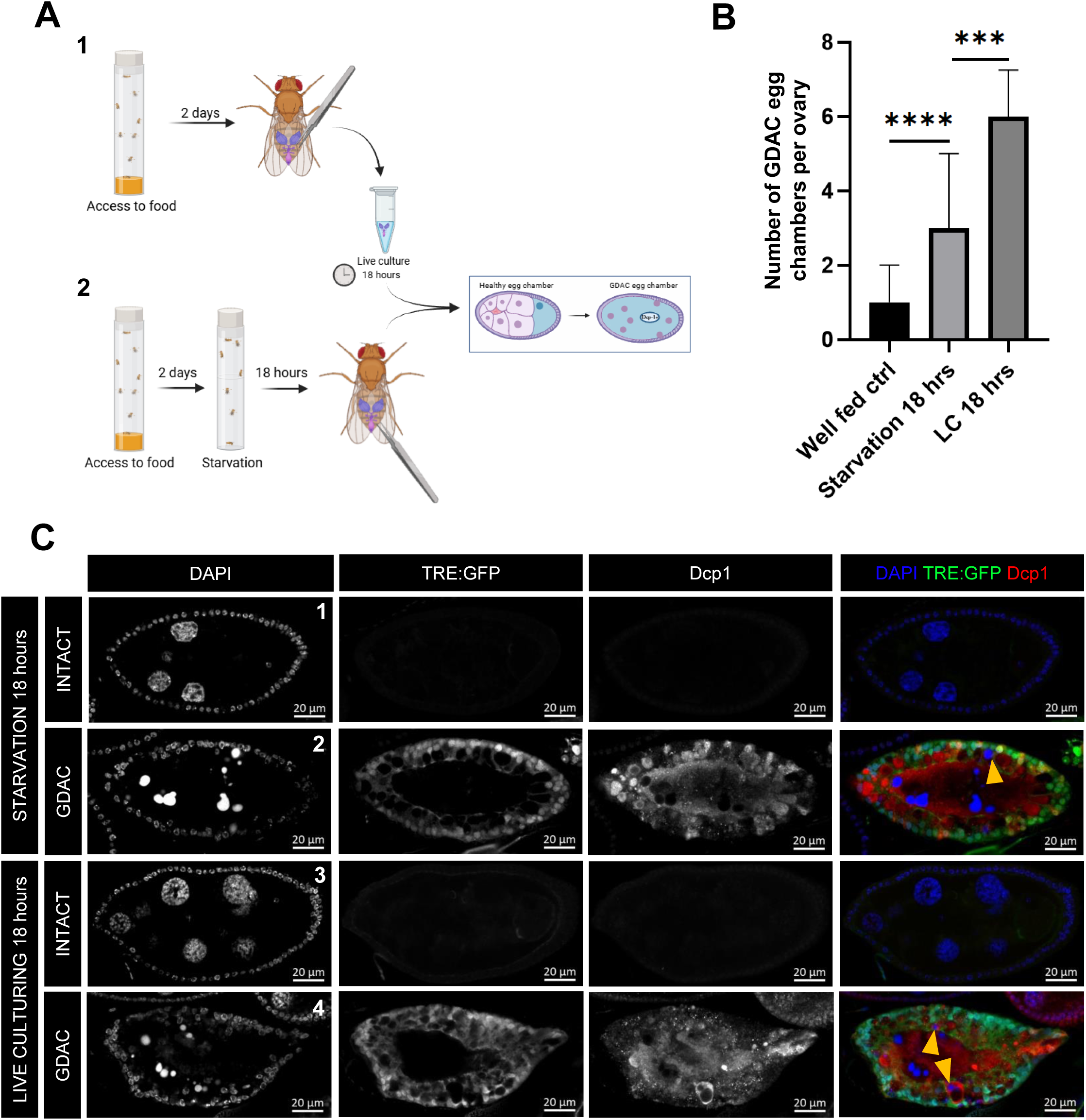
Live culturing as an approach to observe GDAC and FCs to NPPs differentiation. **(A)** Schematic illustration created with BioRender showing the experimental design for inducing and quantifying germline cell death in egg chambers. **A1**: Flies were fed yeast for two days, prior to ovary dissection and culture in *ex vivo* culture medium for 0 (Well-fed control), 6, 12, or 18 hours. **A2**: For starvation treatment, flies were fed yeast for two days, followed by 18 hours of food deprivation on 1% agarose. Ovaries were then dissected and analyzed for GDAC egg chambers using antibody staining against cleaved (active) Dcp-1. **(B)** Bar plots showing the median number of GDAC egg chambers per ovary under each condition, with interquartile range as error bars. Well-fed control (ctrl) (N = 57, median = 1), Live culturing 18 hrs (N = 34, median = 6), Starvation 18 hrs (N = 74, median = 3). P-values obtained from Mann-Whitney test are indicated by *** (p = 0.0004), **** (p ≤ 0.0001). **(C)** Representative confocal images of egg chambers with intact germline cells and GDAC induced by live culturing or starvation. Yellow arrowheads point to fragments of condensed germline nuclei being engulfed by NPPs. DAPI marks nuclei, *TRE:GFP* reports JNK pathway activation, and cleaved (active) Dcp-1 labels dying cells.

In addition to germline cell death, we also confirmed the differentiation of FCs into NPPs under live culturing conditions. Hallmarks of this transition included activation of JNK signaling in NPPs (Costa et al., 2025; Etchegaray et al., 2012; Huang et al., 2025; Serizier et al., 2022), detected using the TPA response element-GFP (*TRE:GFP*) reporter (Fig. 1C4), a well-established readout of JNK pathway activity (Chatterjee and Bohmann, 2012). Additionally, we observed active phagocytosis of germline cell debris by NPPs, further supporting the functional transition of FCs into NPPs (Fig. 1C4). These findings demonstrate that extended *ex vivo* culture is sufficient to trigger GDAC in mid-stage egg chambers, validating this system as a useful model for studying the epithelial-to-NPP transition.

After establishing live culture as a viable model for inducing germline cell death and FC differentiation into NPPs, we performed live imaging on mid-stage *Drosophila* egg chambers to capture the dynamics of this transition in real time (Fig. 2A). Within the first 4 hours of culture, we observed initial expression of the *TRE:GFP* reporter, indicating activation of JNK signaling in FCs. However, at this stage and expression levels, JNK activation alone was insufficient to drive full differentiation, as FCs remained within the epithelial monolayer and did not exhibit overt morphological changes (Fig. 2B, Movie 1).

**Figure 2.**
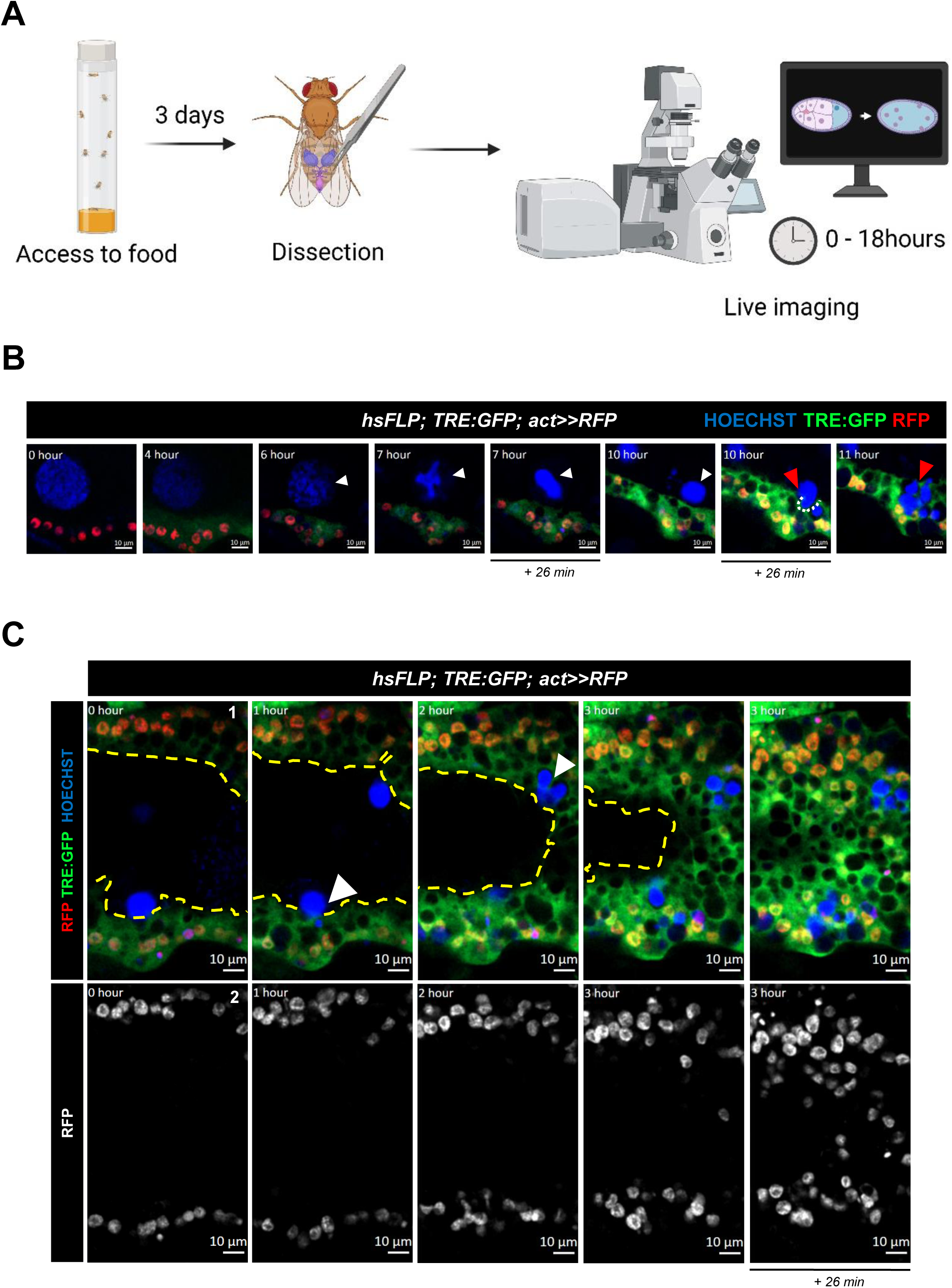
Live imaging of mid-stage egg chambers captures FC-to-NPP differentiation and invasive migration into the germline region. **(A)** Experimental workflow (created with BioRender) showing preparation of *Drosophila* ovaries for live imaging. **(B)** Time-lapse series capturing a germline cell undergoing GDAC and FCs differentiating into NPPs. Hoechst stains nuclei, *TRE:GFP* reports JNK pathway activation and labels the FC cytoplasm (and nucleus), thereby outlining invasive projections, while RFP marks FC nuclei. The white arrowhead highlights DNA condensation in the dying germline nucleus. The red arrowhead shows NPPs forming phagocytic cup-like structures and engulfing the germline remnant. An example of cup-like structure is outlined by a white dashed line **(C)** Time-lapse sequence of a mid-stage egg chamber showing progressive germline nuclear condensation, followed by invasion of the germline region by migratory NPPs and subsequent engulfment of apoptotic germline nuclei over time. Time points are displayed as rounded hours by the ZEISS image software; for consecutive images labeled with the same hour, additional minute annotations were included to indicate temporal progression.

After 7 hours, germline cells began to display nuclear condensation (Fig. 2B, Movie 1), a hallmark of apoptosis, characterized by increased Hoechst fluorescence and progressive chromatin compaction into dense structures resembling those observed in fixed preparations (Fig. 1C2). By approximately 10 hours, nuclear condensation was complete, marking the onset of germline cell death. By this point, FCs had initiated cytoplasmic expansion toward the condensed nuclei and began forming phagocytic cup–like structures to engulf germline debris (Fig. 2B, Movie 1, Fig. S1A).

To visualize changes in individual FC morphology during the transition to NPPs, cells were labeled with *UASp-Lifeact.mGFP* (Fig. S2, Movie 2), a reporter that enables visualization of F-actin dynamics in vivo. At 0 hours, FCs formed a single epithelial layer displaying classical epithelial morphology, characterized by a monolayer of cuboidal cells with well-defined boundaries marked by a cortical actin belt. By 3 hours, coincident with germline nuclear condensation, the cortical actin belt was lost, resulting in diminished distinction between individual epithelial cells and the appearance of small actin-rich protrusions. At 8 hours, NPPs on opposite sides of the egg chamber exhibited pronounced actin-rich protrusions that invaded the germline compartment. By 11 hours, protrusions extending from NPPs on opposing sides established contact, and by 12-13 hours, progressively larger regions of the egg chamber were occupied by extended cytoplasmic protrusions, accompanied by NPP migration toward the germline compartment. During this process, apoptotic germline cell nuclei were engulfed by NPPs.

While previous studies described the FC cytoplasmic enlargement as a morphological change during transition to NPP differentiation (Chasse et al., 2024; Etchegaray et al., 2012; Meehan et al., 2016), using live imaging we could also capture a dynamic and previously unrecognized behavior: FCs acquired invasive and migratory capabilities, actively infiltrating the germline region. Cells located on opposite lateral sides of the egg chamber extended cytoplasmic projections toward one another, as visualized by cytoplasmic TRE:GFP expression. These projections progressively advanced until they met, ultimately occupying the posterior germline territory. The positions of individual migrating cells were independently tracked using nuclear *act>>RFP* labeling, confirming true cellular invasion rather than passive tissue displacement (Fig. 2C, Movie 3, Fig. S1B). Because the invaded region corresponds to the germline territory and lacks both TRE:GFP and RFP expression, restricting these reporters to FCs enables unambiguous tracking of FC invasion into the germline region.

This coordinated migratory invasion reveals a novel, active role for FCs during their transition into NPPs, underscoring the unique advantage of live imaging in capturing transient and dynamic cellular behaviors.

### Hypoxia induces Foxo nuclear localization in germline cells and parallels Foxo activation during *ex vivo* culture

Because *ex vivo* live culturing of egg chambers constitutes an artificial environment, it can expose tissues to multiple stressors, including nutrient deprivation, phototoxic stress during live imaging, and altered oxygen availability. We therefore asked whether hypoxia-responsive stress pathways are activated in live-cultured egg chambers. Foxo (Forkhead box O) is a conserved transcription factor in *Drosophila* that translocates to the nucleus in response to hypoxic and metabolic stress (Elizabeth C. Barretto et al., 2020; Eijkelenboom and Burgering, 2013). Consistent with this, female flies exposed to hypoxic conditions (1% O₂ for 7 h) exhibited robust nuclear localization of Foxo::GFP in germline cells (Fig. S3A, B). Notably, germline cells from live-cultured egg chambers also consistently displayed nuclear Foxo::GFP localization, whereas Foxo::GFP remained largely cytoplasmic in egg chambers from well-fed control flies (Fig. S3C, D). Collectively, these observations indicate that *ex vivo* culture activates stress-responsive signaling pathways similar to those engaged by hypoxia in vivo. However, because Foxo nuclear localization can be induced by multiple forms of cellular stress, these results do not establish hypoxia or metabolic stress as the sole drivers of germline cell death during live culture, but rather provide insight into the stress conditions experienced by egg chambers under *ex vivo* imaging conditions.

### NPP migration and delamination are cell-autonomous behaviors impaired by Notch and JNK signaling disruption

Given the collective migratory behavior exhibited by NPPs, we sought to determine whether this capability was dependent on coordinated interactions with neighboring NPPs or represented an intrinsic, cell-autonomous property of individual FCs. To test this, we generated mosaic egg chambers in which NPPs were positioned adjacent to non-NPP FCs using the heat-shock FLP/FRT system (Golic and Lindquist, 1989) combined with an *act-Gal4* driver (Brand and Perrimon, 1993) flanked by *FRT* sites to drive the expression of *Notch RNAi* (*NIR*). Because Notch signaling is required for JNK activation, polyploidization, and the subsequent differentiation of FCs into NPPs (Huang et al., 2025), *Notch* depletion effectively blocks NPP formation in targeted cells.

We began by comparing fixed mid-stage egg chambers undergoing GDAC (Fig. 3A3, A4) with non-GDAC controls (Fig. 3A1, A2). Hindsight (Hnt) staining confirmed effective *Notch* knockdown in FCs. In non-GDAC egg chambers, both *NIR* and *wild-type* cells retained a monolayer organization (Fig. 3A2). However, under GDAC conditions, *wild-type* NPPs positioned within the germline territory appeared to delaminate from the monolayer formed by adjacent *NIR* clones, which themselves remained in their original epithelial configuration (Fig. 3A3, A4, B, D).

To verify whether germline invasion from active NPPs is due to migration rather than a developmental misplacement, we performed live imaging on egg chambers containing individual NPPs adjacent to non-motile *NIR-*JNK depleted clones. Despite being close by immobile *NIR* cells, the isolated NPP successfully delaminated from the epithelial monolayer and migrated toward the germline territory (Fig. 3C1, Movie 4). To further evaluate this migratory behavior, we quantified the migration velocity of *wild-type* NPPs and *NIR* cells. NPPs exhibited significantly higher migration speeds than *NIR* clones, confirming that motility is a distinct, acquired property of NPPs (Fig. 3E).

**Figure 3.**
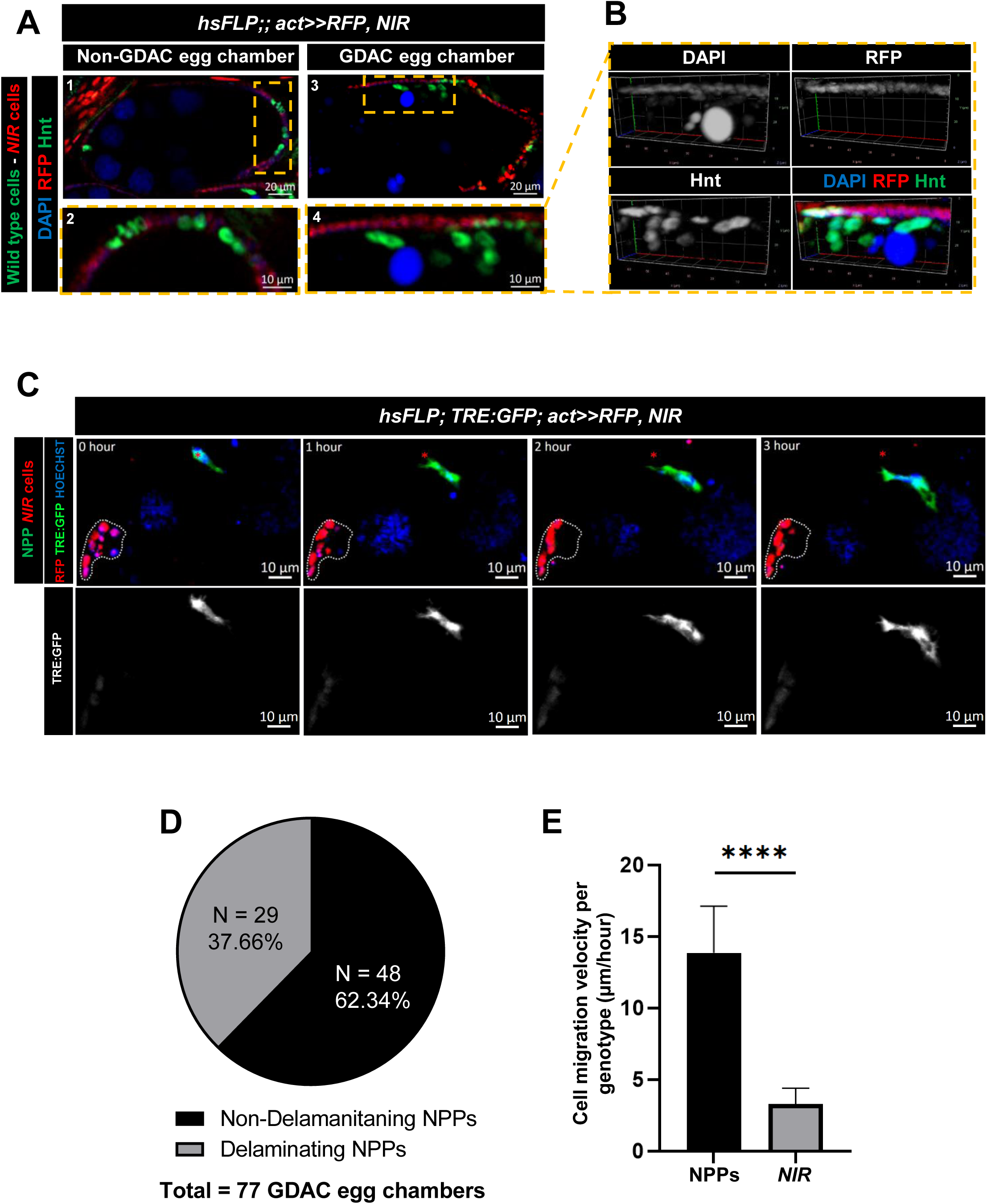
NPP delamination from the follicular epithelium and invasion into the germline region. (**A–B**) All panels show egg chambers containing FLP-out mosaic clones expressing *Notch RNAi* driven by act-Gal4 (*hsFLP;; act>>RFP, NIR*). (**A**) Confocal images of Non-GDAC (**A1–A2**) and GDAC (**A3–A4**) egg chambers highlighting the spatial distribution of *NIR* clones and NPPs. *NIR* clones are marked by RFP expression, while neighboring *wild-type* NPPs are identified by Hindsight (Hnt) staining. Yellow dashed boxes in A2 and A4 indicate the regions magnified in A1 and A3, respectively. (**B**) 3D reconstruction of the region shown in A4. (**C**) Time-lapse sequence of an egg chamber of genotype *hsFLP; TRE:GFP; act>>Gal4, NIR*, showing a delaminated NPP migrating toward the germline region. The red asterisk marks the initial position of the migrating NPP, while white dashed outlines indicate clusters of non-migratory *NIR* cells, which serve as fixed reference points to facilitate visualization of NPP movement. (**D**) Pie chart illustrating the proportion of GDAC egg chambers exhibiting NPP delamination. Out of 77 GDAC egg chambers analyzed, 29 (37.66%) displayed delaminating NPPs, while 48 (62.34%) exhibited non-delaminating NPPs. (**E**) Bar plots showing the median migration velocity (µm/hour) of NPPs (*hsFLP; TRE:GFP; act>>RFP*) versus *NIR* cells (*hsFLP; TRE:GFP; act>>RFP, NIR*), with error bars representing the standard deviation. Migration was quantified for 45 individual cells per group (NPPs: N = 45, mean = 13.86 µm/h; *NIR*: N = 45, mean = 3.28 µm/h). P-value obtained from t-test is indicated by **** (p ≤ 0.0001) above plot. A mid-stage egg chamber area can vary from ∼4,690 to 20,933 µm² (Jia et al., 2016).

Together, these findings demonstrate that delamination and directed migration toward the germline are intrinsic features acquired by FCs during their differentiation into NPPs. These behaviors do not depend on collective coordination with other NPPs and are impaired when *Notch* and JNK signaling are disrupted.

### NPPs engulf and eliminate adjacent *Notch*-depleted FCs

While performing live imaging to assess the migratory and invasive properties of isolated NPPs, we observed dynamic interactions between NPPs and adjacent *NIR* FCs. At 0 hours, a clear boundary was evident between NPPs, marked by TRE:GFP expression, and *NIR* mosaic clones labeled by nuclear RFP (Fig. 4A). By 1 hour, NPPs began extending their cytoplasm toward neighboring *NIR* cells and initiated engulfment of both apoptotic cells exhibiting condensed nuclei and seemingly viable cells with intact nuclei. By 2 hours, *NIR* cells were fully internalized within the NPP cytoplasm.

**Figure 4.**
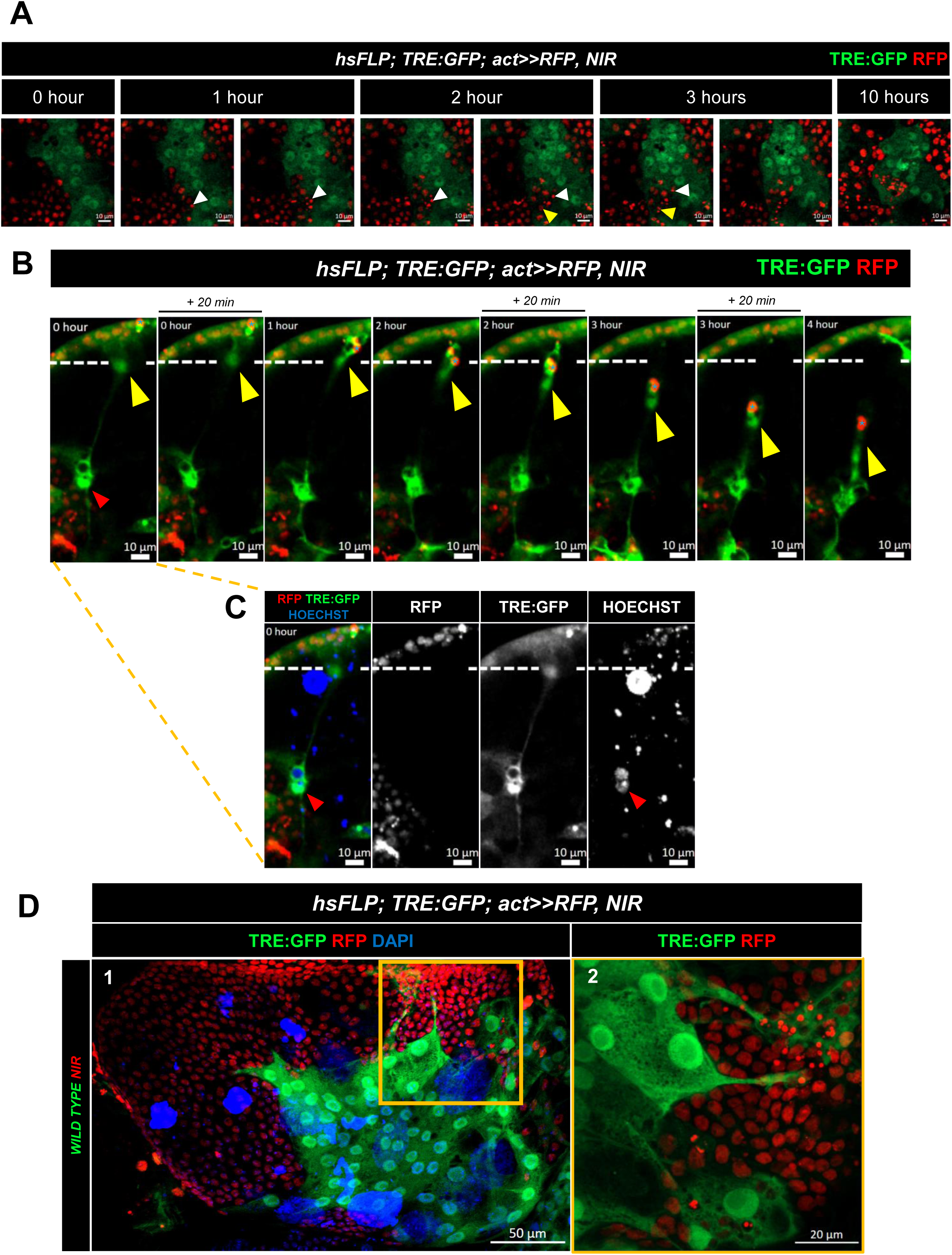
NPPs engulf both apoptotic and live neighboring *NIR* cells. (**A**, **B**) Time-lapse sequences showing dynamic interactions between *TRE:GFP*-positive NPPs and *NIR* clones. NPPs engage with both apoptotic *NIR* cells exhibiting nuclear condensation and viable *NIR* cells with intact nuclei. (**A**) White arrowhead points to an apoptotic *NIR* (condensed nucleus) being engulfed gradually by NPPs, yellow arrowhead points to a viable *NIR* cell (non-condensed nucleus) that is engulfed and subsequently degraded by NPPs. (**B**) Yellow arrowheads indicate a mobile TRE:GFP-positive cytoplasmic projection that retracts over time following the capture of an apoptotic *NIR* cell (blue asterisk), which exhibits a condensed nucleus labeled by RFP. The projection retracts toward the cell of origin, marked by a red arrowhead. The white dashed line denotes the initial position of the cytoplasmic projection. Time points are displayed in hours as provided by the ZEISS software; for consecutive images labeled with the same hour, additional minute annotations were included to indicate temporal progression. (**C**) Corresponding 0-hour image from panel (**B**) shown as separated channels (RFP, TRE:GFP, and Hoechst). The red arrowhead indicates the nucleus of a single NPP. (**D1**) Confocal image of a fixed GDAC egg chamber containing a mosaic of *NIR* clones (marked by RFP) and *wild-type* NPPs marked by *TRE:GFP*, indicating JNK activation. (**D2**) Higher magnification of the boxed region in **D1** highlighting NPPs extending cytoplasmic projections and phagocytosis of apoptotic *NIR* cells.

Following internalization, nuclear condensation was observed in previously viable engulfed *NIR* nuclei, suggesting that engulfment alone is sufficient to induce cell death. Between 3 and 10 hours, additional *NIR* cells were progressively engulfed, and internalized nuclei underwent fragmentation into smaller RFP-positive nuclear remnants, indicating that engulfment precedes and drives nuclear fragmentation (Fig. 4A, Movie 5, Fig. S1C).

In addition, NPPs exhibited pronounced cell body elongation and target capture behaviors. During these events, individual TRE:GFP-expressing NPP underwent dynamic morphological remodeling, extending long cytoplasmic protrusions. This extension enabled the capture of apoptotic *NIR* cells, as indicated by the presence of condensed nuclei. Time-lapse imaging further reveals that these projections retract after target engagement, suggesting an active mechanism for long-range capture and internalization of apoptotic debris (Fig. 4B, C; Movie 6).

To investigate this interaction in more detail, we analyzed fixed GDAC egg chambers containing mixed populations of NPPs and *NIR* clones. Remarkably, we found that *NIR* cells underwent apoptosis, as evidenced by condensed nuclei, when located adjacent to *wild-type* NPPs. Consistent with the dynamic behaviors observed in live imaging (Fig. 4B, C; Movie 6), these NPPs responded by extending pseudopodia-like projections toward the apoptotic *NIR* cells, engulfing them even from a distance (Fig. 4D1, D2).

Overall, these findings reveal a versatile phagocytic behavior acquired by epithelial FCs upon NPP differentiation. This includes the ability to eliminate not only apoptotic cells but also seemingly viable neighboring epithelial cells, uncovering previously unrecognized strategies by which NPPs ensure effective clearance of surrounding cell populations.

### Germline cell death is accompanied by actin cytoskeleton remodeling and perinuclear ring formation

Given our ability to use live imaging to visualize, in real time, the progressive death of germline cells and the associated nuclear condensation, we concluded this study by examining dynamic changes in the actin cytoskeleton during this process. Since the mechanisms regulating germline cell death in mid-stage egg chambers remain poorly understood, we anticipated that this analysis would offer novel insights into the structural events accompanying germline cell apoptosis. To monitor actin dynamics specifically in the germline, we expressed *UASp-Lifeact.mGFP* using the germline-specific *CG6325Gal4* driver, which delineates the cellular membrane of germline cells and allows real-time visualization of structural changes.

Live imaging revealed a progressive reorganization of actin filaments during germline cell death. This process began with a gradual loss of cortical actin organization between germline cells, preceding the onset of nuclear condensation. (Fig. 5A, 60 min), followed by a pronounced accumulation of actin filaments around the nucleus (Fig. 5A, 89 min). With continued nuclear compaction, perinuclear actin enrichment intensified, forming a distinct ring-like structure that progressively decreased in diameter in parallel with nuclear shrinkage (Fig. 5A, 2 h; Movie 7).

**Figure 5.**
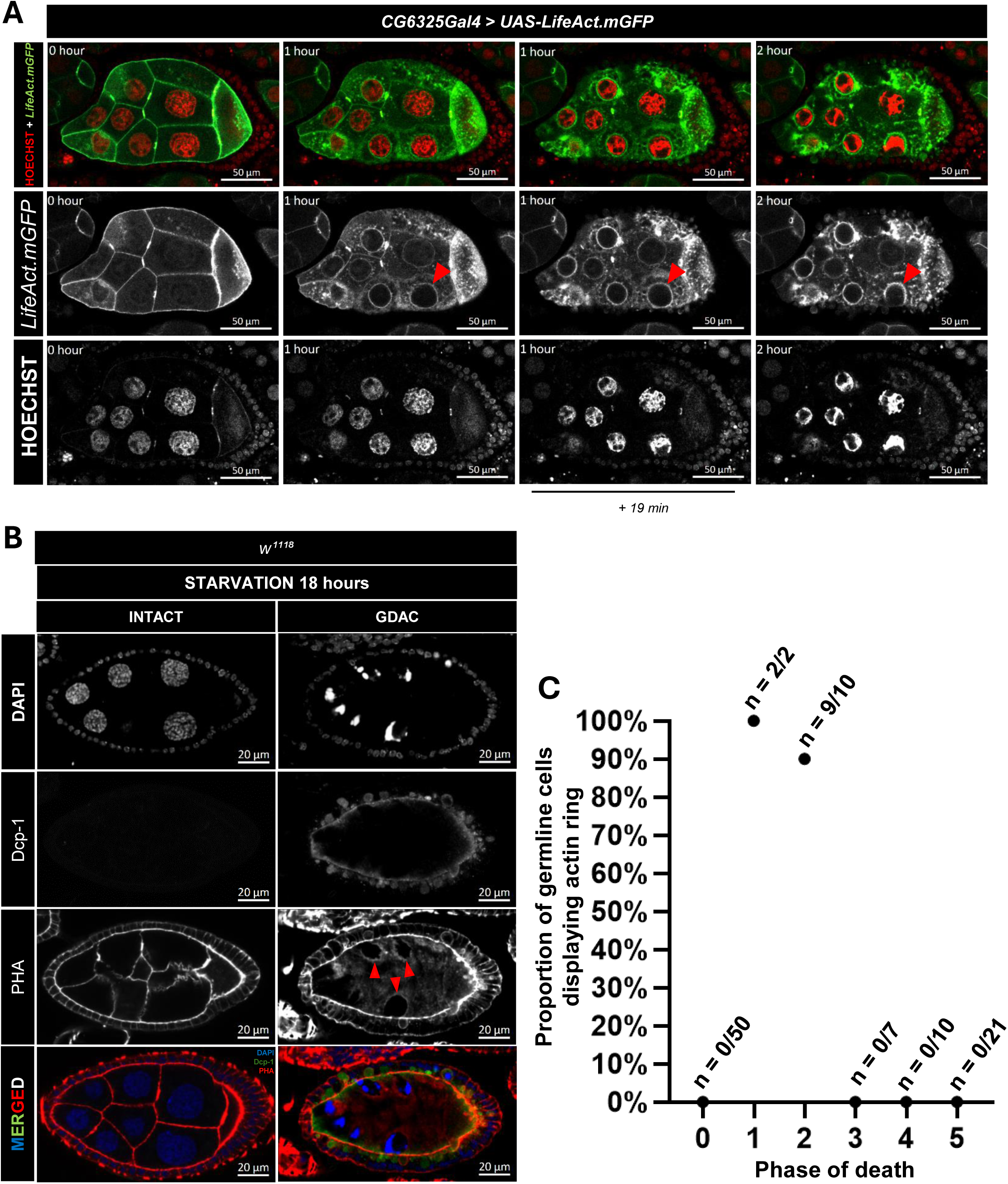
Actin cytoskeleton remodeling during germline cell death. (**A**) Time-lapse imaging of a mid-stage egg chamber undergoing germline cell death as shown by progressive nuclear condensation (Hoechst) and changes in actin cytoskeletal organization as revealed by *UASp-Lifeact.mGFP* specifically expressed in germline cells. Red arrowheads indicate perinuclear actin ring formation detected by Lifeact.mGFP in an egg chamber undergoing GDAC. Time 0 corresponds to an egg chamber imaged 6.5 hours prior to the onset of nuclear condensation. Time points are displayed in hours as provided by the ZEISS software; for consecutive images labeled with the same hour, additional minute annotations were included to indicate temporal progression. (**B**) Confocal images of egg chambers showing differences in actin cytoskeletal organization between fixed GDAC egg chambers containing dying germline cells and Non-GDAC egg chambers displaying intact germline cells. Red arrowheads indicate perinuclear actin rings detected by PHA staining in a GDAC egg chamber. **(C)** Dot plot showing the proportion of egg chambers exhibiting the actin ring phenotype in germline cells, categorized by phase of germline cell death. Egg chambers with morphologically intact germline cells were classified as phase 0 (N = 50). Egg chambers showing signs of germline cell death were divided into phases 1–5 (total N = 50). The percentage of actin ring–positive egg chambers in each phase is as follows: phase 0 (0/50; 0%), phase 1 (2/2; 100%), phase 2 (9/10; 90%), phase 3 (0/7; 0%), phase 4 (0/10; 0%), and phase 5 (0/21; 0%)

Notably, similar actin ring structures were observed in dying egg chambers from starved flies (Fig. 5B), suggesting that this phenomenon is not an artifact of *ex vivo* culturing, but rather a physiological event. These structures were predominantly observed during the early phases of germline apoptosis, specifically phases 1 and 2, when chromatin is disordered (phase 1) and displays condensation (phase 2) as previously described (Etchegaray et al., 2012). In contrast, actin rings were absent in germline cells from phase 0 (healthy), where egg chambers display dispersed chromatin, and from more advanced apoptotic phases (phases 3–5), during which nuclei become highly condensed and fragmented, indicating that this is a transient and early germline-cell-death-specific feature (Fig. 5C).

To understand the transient nature of actin ring formation upon full condensation of germline cell nucleus, we extended live imaging beyond 8 hours (Fig. S4A, Movie 8). Over time, the perinuclear actin rings gradually decreased in diameter and intensity as nuclear condensation advanced (Fig. S4A, 9-14 hours; Fig. S4B). Once the nucleus reached full compaction, the actin ring structure demonstrated signs of disappearance and morphological change (Fig. S4A, 14 hour). This temporal pattern suggests that the perinuclear actin ring is a transient, phase-specific structure that emerges during early apoptosis and is lost as germline cells progress to full nuclear condensation.

Although not always in direct contact with nuclear chromatin, these actin structures were consistently positioned around condensing nuclei, suggesting a potential mechanical role. We hypothesize that the actin ring may exert compressive forces that facilitate nuclear compaction, either by directly squeezing the nucleus or by influencing other nuclear envelope components. This points to a possible cytoskeletal mechanism contributing to programmed germline cell death.

## DISCUSSION

In this study, we used live imaging of mid-stage egg chambers to capture the dynamic transition of epithelial FCs into NPPs. Given that *ex vivo* culture of live tissue can introduce multiple apoptotic stressors, including nutrient deprivation, hypoxia, and laser exposure (Icha et al., 2017; Sheard and Cox, 2021; Shimada et al., 2011), this experimental setup enabled us to induce GDAC and capture the NPP transformation.

*Ex vivo* culture is likely to expose egg chambers to multiple stressors, including altered oxygen and nutrient availability as well as mechanical stress. Our observation that Foxo is expressed in the nucleus of both live-cultured egg chambers and hypoxia-treated egg chambers suggests that overlapping stress-responsive pathways are engaged under these conditions. Foxo nuclear translocation has been linked to cellular metabolic stress responses and has also been implicated as a mediator of stress-induced apoptotic programs (Elizabeth C Barretto et al., 2020; Jünger et al., 2003; Luo et al., 2007), potentially contributing to sensitization of germline cells to death. However, because Foxo nuclear localization can be induced by diverse forms of cellular stress (Elizabeth C Barretto et al., 2020; Jünger et al., 2003; Luo et al., 2007), it cannot be used to distinguish among nutrient deprivation, hypoxia, or other stress modalities. Future studies directly monitoring oxygen availability in culture media and genetically manipulating hypoxia-responsive signaling pathways will be required to determine whether hypoxia plays a causal role in driving germline cell death in live-cultured egg chambers.

While these stress-responsive stimuli likely created a permissive context for germline cell death, our primary focus was to use this approach to uncover cytoplasmic remodeling and the acquisition of migratory behavior by FCs as they transition into NPPs, equipping them to perform phagocytic functions under diverse conditions. These findings broaden our understanding of epithelial plasticity and the functional versatility of NPPs in vivo.

Our findings reveal that JNK-activated FCs not only undergo cytoplasmic expansion, as previously documented in fixed imaging, but also display coordinated migratory behavior. These cells actively and progressively invade the degenerating germline territory until it is fully occupied. Strikingly, the ability of FCs to extend cytoplasm while maintaining epithelial sheet integrity and cohesion resembles the collective invasive behavior observed in epithelial cells during wound healing and cancer invasion (Friedl and Gilmour, 2009). This establishes FC-derived NPPs as a model to study epithelial coordination during tissue remodeling.

Interestingly, the migratory and phagocytic capabilities displayed by NPPs show partial resemblance to other somatic cell populations in *Drosophila*. Similar to border cells (Dai and Montell, 2016; Miao et al., 2022; Prasad et al., 2015; Prasad and Montell, 2007), NPPs exhibit collective cell migration involving coordinated epithelial movement. However, whereas NPP migration is triggered in response to germline cell death and facilitates the invasion of the germline region to engulf dying cells and cellular debris, border cell migration occurs strictly within a developmental context and is not associated with phagocytic activity.

A closer functional parallel to NPPs can be found in somatic cyst cells in the testis, which actively eliminate germline cell progenitors via phagoptosis (Zohar-Fux et al., 2022). Notably, unlike NPPs, cyst cells largely remain stationary and already encapsulate germ cells, eliminating the need for long-range or invasive migration. Thus, ovarian NPPs uniquely integrate epithelial collective migration with non-professional phagocytic activity, establishing FC–derived NPPs as a distinct model of epithelial phagocytes.

Even when placed adjacent to *NIR* non-NPP clones, *wild-type* NPPs retained their ability to delaminate from the epithelial monolayer, underscoring the cell-autonomous nature of their migratory and phagocytic program. This autonomous behavior mirrors that of professional phagocytes such as macrophages, which must migrate independently from the vasculature, traversing endothelial barriers, to reach inflamed tissues where pathogens or dying cells are present. Like NPPs, macrophages execute this migratory and phagocytic response without requiring multicellular coordination, relying instead on intrinsic signaling pathways and cytoskeletal dynamics (Gao et al., 2021).

Beyond engulfing dying germline cells, NPPs also extended cytoplasmic projections resembling macrophage lamellipodia and formed phagocytic cup–like structures to target apoptotic neighboring cells (Jaumouillé and Waterman, 2020). More remarkably, we observed NPPs engulfing *NIR* cells with intact nuclei. These internalized cells subsequently underwent nuclear fragmentation, indicating that cell degradation was induced after engulfment. This phenomenon resembles the definition of phagoptosis, a process by which viable cells are phagocytosed and subsequently eliminated, previously described in macrophages (Brown and Neher, 2012), and during normal germline cell death in late-stage *Drosophila* ovarian egg chambers (Yalonetskaya et al., 2018).

Additionally, by driving germline-specific expression of Actin-GFP, we found that germline nuclear condensation is accompanied by actin cytoskeletal remodeling and preceded by a loss of cortical actin organization between germline cells. This loss of cortical actin suggests disruption of germline cell boundaries; however, as actin is an indirect marker, it does not directly report plasma membrane integrity. Therefore, we focus our interpretation on changes in actin organization rather than definitive membrane breakdown. We also captured the formation of dynamic perinuclear actin rings that progressively contract as germline nuclei undergo compaction. Similar actin-rich structures have been reported in mouse pre-granulosa cells (Niu and Spradling, 2022), suggesting that perinuclear actin remodeling may represent a conserved mechanism associated with nuclear condensation during apoptosis across species.

Previous live-imaging studies of the *Drosophila* ovary have primarily been designed to preserve tissue homeostasis and maintain germline viability in order to visualize developmental processes in the germarium and across different egg-chamber stages (Airoldi et al., 2011; Bianco et al., 2007; Cai et al., 2014; Cetera et al., 2014; Dorman et al., 2004; Fichelson et al., 2009; Forrest and Gavis, 2003; Gilliland et al., 2007; He et al., 2010; Jackson et al., 2023; McLean and Cooley, 2013; Morris and Spradling, 2011; Osterfield et al., 2013; Peters and Berg, 2016; Prasad et al., 2007; Spracklen et al., 2014; Tekotte et al., 2007; Tootle and Spradling, 2008; Wang and Hazelrigg, 1994; Wang et al., 2010; Zhao et al., 2012). Although these studies typically present images of morphologically healthy egg chambers, their emphasis on developmental events does not necessarily imply that germline cell death does not occur during *ex vivo* culture; rather, such events were outside the scope of investigation and often beyond the temporal window analyzed and therefore were not chosen to be displayed on the studies. In contrast, the defining feature of our study is the use of live imaging specifically to follow the progression of germline cell death and the subsequent differentiation of FCs into NPPs. A key distinction of our protocol is the extended culture duration, which allows these events to unfold. Whereas most previous ovary live-imaging studies limit culture duration to approximately 11 hours or less, we extended imaging to up to 20 hours, enabling reproducible observation of germline apoptosis and epithelial phagocytic behaviors.

Because prolonged *ex vivo* culture raises the concern of artifactual effects on cell behavior, we validated all major phenotypes observed by live imaging using corresponding fixed samples that were not subjected to live culture. In addition, although Hoechst labeling has been reported to induce cytotoxicity during long-term time-lapse imaging due to phototoxicity from repeated excitation, we minimized this risk by restricting Hoechst exposure to a brief pre-incubation (1.5 h) followed by complete dye removal prior to imaging, as well as by reducing 405-nm laser intensity.

Importantly, our live-culture medium is comparable to that used in established ovary live-imaging protocols and consists of Schneider’s medium supplemented with penicillin–streptomycin, fetal bovine serum, and insulin (Airoldi et al., 2011; Bianco et al., 2007; Cai et al., 2014; Cetera et al., 2014; He et al., 2010; McLean and Cooley, 2013; Montell et al., 1991; Morris and Spradling, 2011; Osterfield et al., 2013; Prasad and Montell, 2007; Wang et al., 2010). These considerations indicate that the primary distinction of our approach lies not in the culture conditions themselves, but in the deliberate choice to image egg chambers over extended periods to capture germline cell death and the dynamic epithelial transition to NPPs, processes that are missed in homeostasis-focused imaging paradigms.

Taken together, these findings reveal a broad spectrum of previously unrecognized capabilities acquired by epithelial FCs during their transition into NPPs, driven by activation of JNK signaling. These behaviors, ranging from autonomous migration and phagocytosis to cell death progression, occur dynamically and with rapid temporal resolution. Live imaging thus uncovers that this transition is not merely a passive response to neighboring cell death but a highly active, coordinated, and multifaceted process. These insights not only advance our understanding of epithelial plasticity in *Drosophila* but may also inform studies of epithelial-mediated clearance, tissue remodeling, and cell competition in higher organisms.

## METHODS

### Fly Stocks and Husbandry

Flies were maintained under standard laboratory conditions at 25 °C and fed with yeast for three days prior to dissection. For starvation experiments, flies were fed yeast for two days, followed by 18 hours of food deprivation, as described in previous protocols (Bandyadka et al., 2025; Lebo et al., 2021) by transferring them to vials containing 1% agar (Apex, Cat# 66-104) dissolved in distilled water.

Mosaic clones were generated using a heat-shock–inducible FLP/FRT system (*hsFLP*) combined with an actin promoter–driven FLP-out cassette (*act>>RFP*). Upon heat shock, FLP-mediated recombination excises a transcriptional STOP cassette, enabling Gal4 expression under the control of the actin promoter in recombined cells. This, in turn, drives the expression of UAS-linked transgenes, including *Notch RNAi* (*UAS-NIR*), specifically within mosaic clones.

To induce partial mosaicism, flies were subjected to a 15-minute heat shock at 37 °C, while a 40-minute heat shock at the same temperature was used to induce near-complete mosaicism, in which the vast majority of cells underwent FLP-mediated recombination and expressed Gal4-driven transgenes.

We thank the Bloomington *Drosophila* Stock Center for providing the lines used in this study: *NIR* (#33616), *CG6325Gal4* (#54593), *UASpAct-GFP* (#58718) and Dr. Bohmann for providing *TRE:GFP* (Chatterjee and Bohmann, 2012). In the *hsFLP; act>>RFP* system, RFP labels FC nuclei, enabling clear discrimination of cell position, while *TRE:GFP* labels both the nucleus and cytoplasm, allowing visualization of invasive cytoplasmic projections.

### Hypoxia exposure

Larvae of the indicated genotypes were raised on standard food at 25 °C from the first instar stage until adulthood. Five-day-old adult flies were maintained under normoxic conditions in vials exposed to ambient air. For hypoxia experiments, mated female adults were grouped at 15 flies per vial and exposed to a hypoxic environment (1% O₂) for 7 h. Following hypoxic treatment, vials were removed from the hypoxia chamber, and ovaries were immediately dissected, fixed, and processed for immunostaining.

### Live Imaging Sample Preparation

To capture the dynamic differentiation of FCs into NPPs, we cultured mid-stage *Drosophila* egg chambers in *ex vivo* culture medium. Ovaries were dissected in pre-filtered Shields and Sang M3 insect medium (SD media) supplemented with Hoechst 33342 (20 mM stock solution; Thermo Scientific, Prod. 62249) at a 1:1000 dilution to stain nuclei. Dissected ovaries were gently nutated in this solution for 1 hour and 30 minutes at room temperature.

Following staining, ovaries were transferred onto Cell Imaging Dish with glass bottom (Eppendorf AG, 145um, 35 x 10 mm, Cat. No.: 0030740009). Excess SD medium containing Hoechst was carefully removed to reduce background fluorescence. A small volume of 2% low-melting-point agarose (IBI Scientific, IB70051) prepared in PBS, preheated to 87 °C and cooled to 37 °C, was added to cover the samples and immobilize the egg chambers. To expedite agarose solidification and minimize the time the samples remained without nutrients, the culture disc was placed in a 4 °C refrigerator for 1 minute and 20 seconds.

To further stabilize the samples during imaging, three layers of sterile Kimwipe paper (Kimberly-Clark, Kimwipes Kimtech Science) were gently placed on top, followed by the addition of 250 μL of live imaging culture medium. The medium composition was adapted from a previously established protocol for long-term live imaging of *Drosophila* larval imaginal discs (Tsao et al., 2016) and consisted of 95% Schneider’s Insect Medium (Gen Clone, REF. 21720-024), 2% Fetal Bovine Serum (FBS) (Avantor, Product No. 89510-186), 2% Insulin (Sigma, SLCB9851), and 1% Penicillin–Streptomycin (Gibco, REF. 15140-122).

A custom silicone spacer was fabricated using the SYLGARD™ 184 Silicone Elastomer Kit (GMID: 04019862) by mixing 90% of Part A with 10% of Part B and pouring it into a cell imaging dish. After curing for 48 hours, the mold was used to gently secure the sample and ensure consistent pressure during imaging. To prevent dehydration, the entire culture dish was sealed with Parafilm (Bemis, Laboratory Film 2 IN. X 250 FT.).

### Confocal Imaging

Live imaging was performed using a Zeiss LSM 980 confocal microscope, while images used for quantification were acquired on a Zeiss LSM 900. Both systems were operated using consistent imaging parameters: line averaging mode of 4, scan speed of 8, and multi-position acquisition via tile scanning. The master gain was set between 750 and 800, and laser power was minimized to the lowest level necessary for fluorescence detection to reduce phototoxicity and preserve tissue integrity.

Representative images were captured using a Zeiss LSM 800 confocal microscope. For most samples, the master gain was set between 780 and 800, and laser intensity was carefully optimized to prevent photobleaching. A line averaging mode of 8 and a scan speed of 6 were typically used to achieve high-resolution imaging. All image acquisition, processing, and analysis were performed using ZEN Blue software (Zeiss).

### Immunohistochemistry

Ovaries were dissected at room temperature in 1× PBS and fixed in 4% paraformaldehyde (PFA) for 15 minutes. After fixation, samples were washed for 30 minutes in 1× PBT (PBS containing 0.2% Triton X-100) to permeabilize the tissue. Ovaries were then incubated overnight at 4 °C with the primary antibody diluted in PBT.

Following primary antibody incubation, samples were washed in 1× PBT for 30 minutes and incubated with the secondary antibody, also diluted in PBT, either overnight at 4 °C or for 2 hours at room temperature. After secondary incubation, samples were washed again in 1× PBT for 30 minutes, then counterstained with DAPI (Invitrogen, 1 μg/mL) to label nuclei. For actin staining, Phalloidin Fluor 633 (Invitrogen, A22284) was diluted 1:50 in the DAPI solution.

Finally, samples were mounted in 80% glycerol mounting medium and placed onto confocal-compatible microscope slides for imaging.

The primary antibodies used were mouse anti-Armadillo (N27A1, 1:40), anti-Hindsight (1G9, 1:15) obtained from the Developmental Studies Hybridoma Bank (DSHB) and rabbit anti-Dcp-1 (Asp216) from Cell Signaling. Secondary antibodies included Alexa Fluor 488, 546, and 633 (1:400, Molecular Probes).

### Cellular Migration Velocity

Cellular migration velocity was calculated by measuring the displacement of individual cells between time points using the line measurement tool in ZEISS ZEN software. Specifically, a line was drawn from the center of the cell nucleus at the initial time point to its position at the subsequent time point. The measured distance (in micrometers, µm) was divided by the elapsed time (in hours) to determine migration velocity, following established methods (Kanazawa et al., 2010; Voon et al., 2022). A total of 15 randomly selected single cells were analyzed per group across three independent replicates for both NPPs and *NIR* cells. All measurements were tested for normality, and outliers were assessed using standard detection methods. The datasets passed the normality test, and no outliers were identified for removal. Replicates within each group were compared using a t-test, which revealed no significant differences (p > 0.05), supporting the pooling of the data. The final dataset consisted of 45 measurements per group, and statistical comparison between NPPs and *NIR* cells was performed using an unpaired t-test with Welch’s correction.

### Quantitative analysis

Images used for quantification were acquired using a Zeiss LSM 900 confocal microscope. For each experimental condition, well-fed, live culturing at 6, 12, and 18 hours, and 18-hour starvation, a minimum of 15 age-matched female flies were analyzed. All flies were maintained under standardized conditions prior to dissection and processed using the same dissection and immunostaining protocol.

Egg chambers were classified as undergoing GDAC if they exhibited both nuclear condensation in the germline and staining for cleaved (active) Dcp-1. GDAC-positive egg chambers were quantified using the “Custom Graphics” event marking tool in the ZEN Blue microscope software (Zeiss).

Quantification data were analyzed using GraphPad Prism (version 9.5.1 for Windows, GraphPad Software, Boston, MA, USA, www.graphpad.com). Prior to statistical analysis, identified outliers were excluded. Data distributions were assessed for normality using Prism’s built-in normality tests. When datasets met parametric assumptions, comparisons between two groups were performed using Welch’s *t*-test. When normality was not satisfied, the non-parametric Mann–Whitney U test was applied.

For fluorescence intensity analyses, raw GFP and DAPI intensities were first evaluated independently for normality and intergroup differences. DAPI intensity did not show statistically significant differences between groups and was therefore used as a reference for normalization. GFP intensity values were normalized to DAPI by calculating the GFP/DAPI ratio for each sample. Normality of the resulting normalized datasets was reassessed prior to statistical comparison between groups using the appropriate parametric or non-parametric test. Statistical significance was defined as p < 0.05.

Graphs were generated using GraphPad Prism. For non-parametric analyses (Mann–Whitney U test), data are presented as median with interquartile range (IQR). For parametric analyses (Welch’s *t*-test), data are presented as mean ± standard error of the mean (SEM). Corresponding *p*-values are indicated on the graphs.

### Scientific writing and proofreading

The authors acknowledge the use of ChatGPT 4o (OpenAI) as a tool for initial grammar review and improving readability during manuscript preparation. They also thank Elinor A. Romaguera for her final grammar review and contributions to the clarity of the text. The authors take full responsibility for the scientific content and conclusions of this study.

## Acknowledgments

We thank the Bloomington *Drosophila* Stock Center for providing fly lines and the Developmental Studies Hybridoma Bank for providing antibodies.

