## Supplementary figures and images for "Live Imaging of Epithelial Phagocyte Differentiation in the *Drosophila* Ovary Reveals Transient and Novel Behaviors"

### Supplemental Figure 1

#### Supplementary figure 1

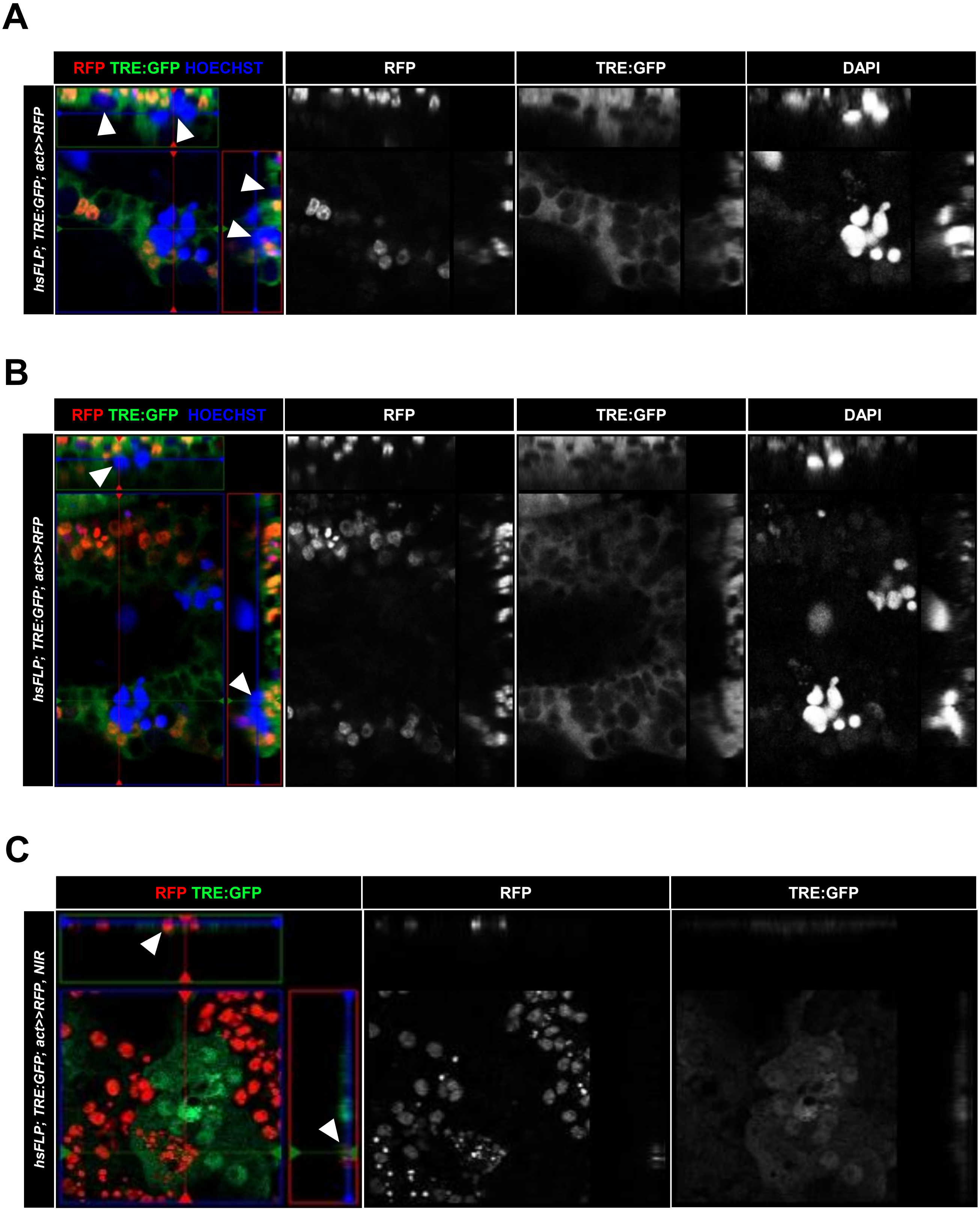

### Supplemental Figure 2

#### Supplementary figure 2

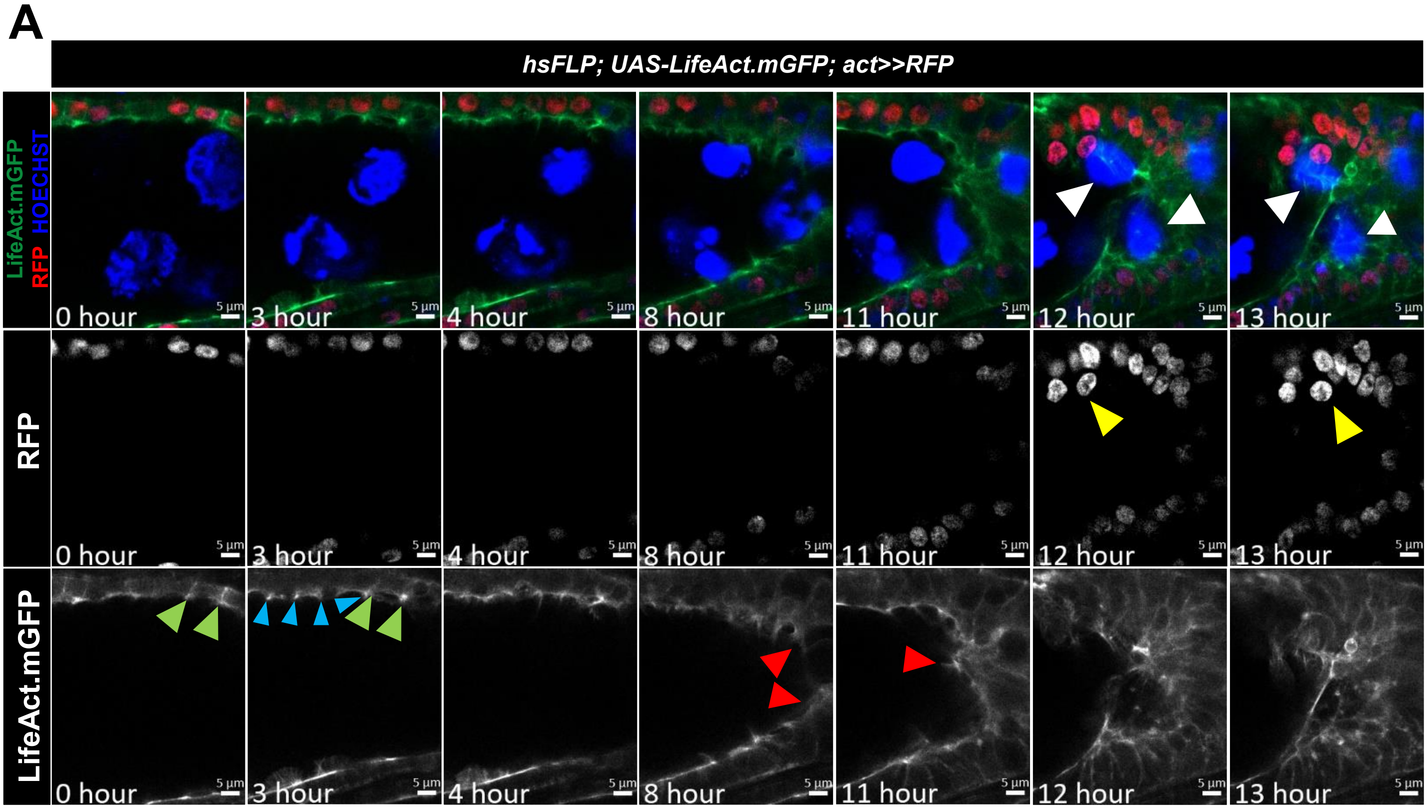

### Supplemental Figure 3

#### Supplementary figure 3

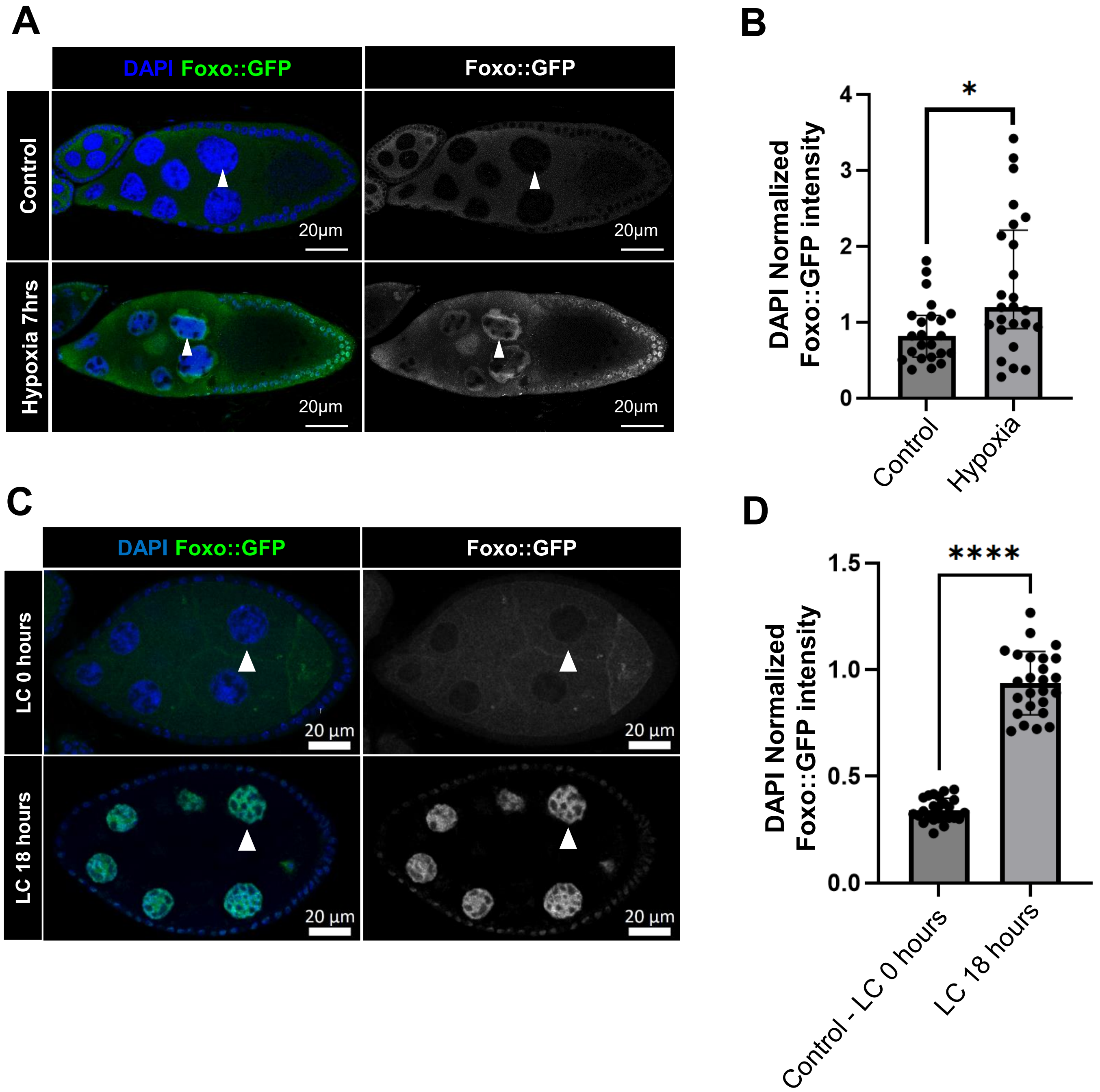

### Supplemental Figure 4

#### Supplementary figure 4

A

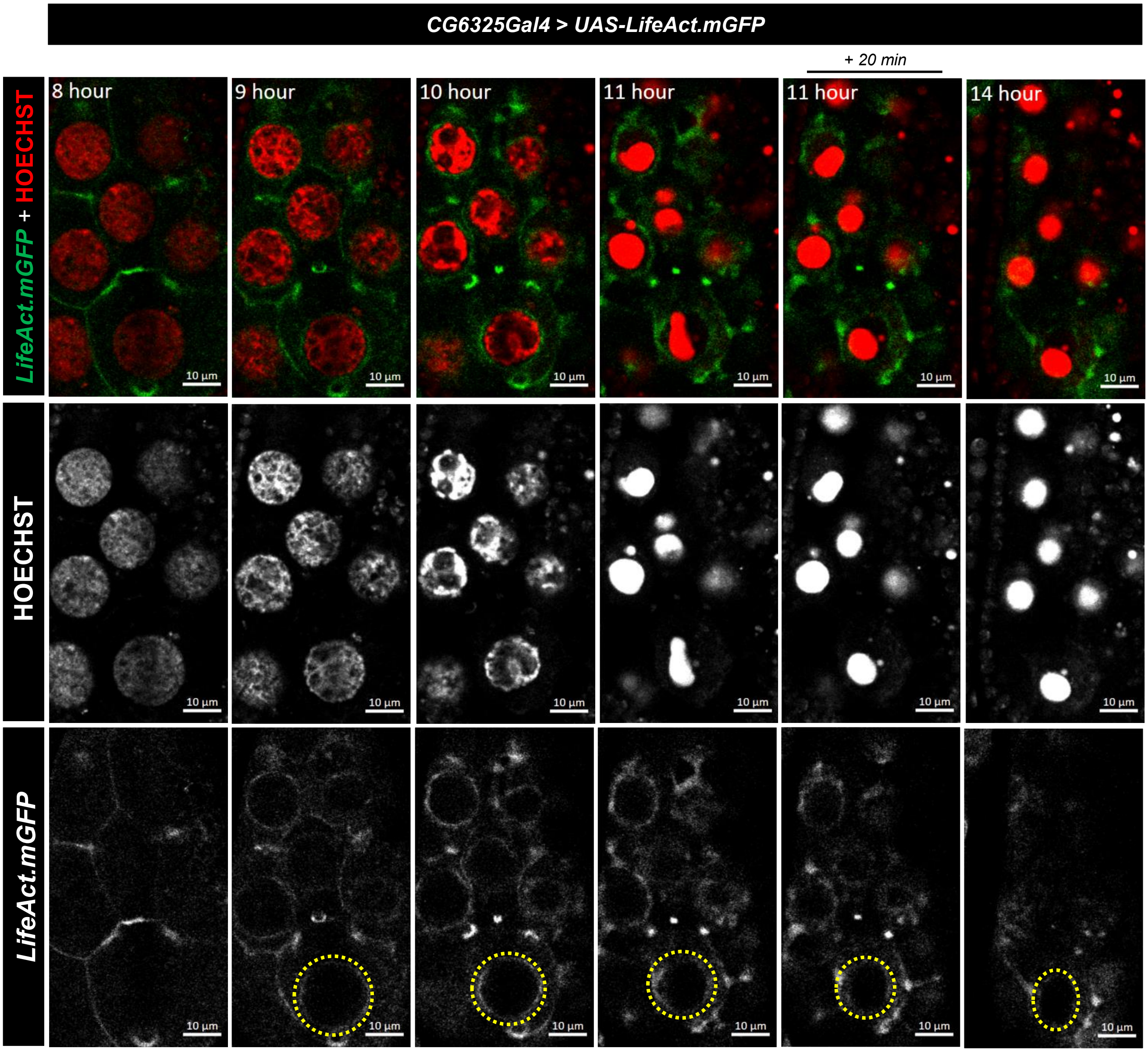


B

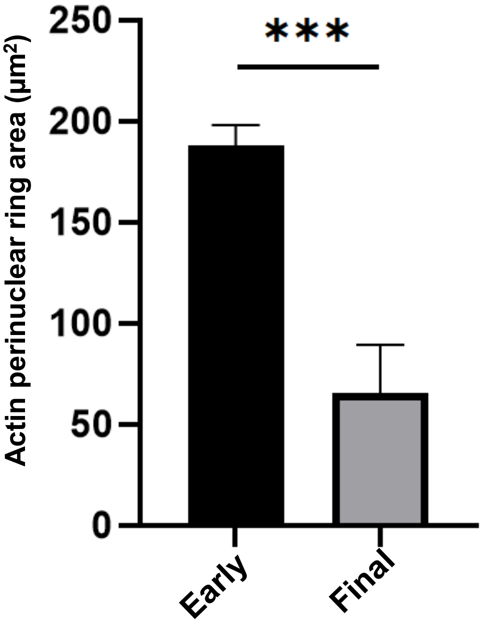
